# RFDTI: Using Rotation Forest with Feature Weighted for Drug-Target Interaction Prediction from Drug Molecular Structure and Protein Sequence

**DOI:** 10.1101/2020.01.06.895755

**Authors:** Lei Wang, Zhu-Hong You, Li-Ping Li, Xin Yan

**Affiliations:** College of Information Science and Engineering, Zaozhuang University, Zaozhuang, 277160, China; Xinjiang Technical Institutes of Physics and Chemistry, Chinese Academy of Sciences, Urumqi, 830011, China; School of Foreign Languages, Zaozhuang University, Zaozhuang, 277160, China

**Keywords:** drug-target interaction, rotation forest, pseudo position-specific score matrix

## Abstract

The identification and prediction of Drug-Target Interactions (DTIs) is the basis for screening drug candidates, which plays a vital role in the development of innovative drugs. However, due to the time-consuming and high cost constraints of biological experimental methods, traditional drug target identification technologies are often difficult to develop on a large scale. Therefore, *in silico* methods are urgently needed to predict drug-target interactions in a genome-wide manner. In this article, we design a new *in silico* approach, named RFDTI to predict the DTIs combine Feature weighted Rotation Forest (FwRF) classifier with protein amino acids information. This model has two outstanding advantages: a) using the fusion data of protein sequence and drug molecular fingerprint, which can fully carry information; b) using the classifier with feature selection ability, which can effectively remove noise information and improve prediction performance. More specifically, we first use Position-Specific Score Matrix (PSSM) to numerically convert protein sequences and utilize Pseudo Position-Specific Score Matrix (PsePSSM) to extract their features. Then a unified digital descriptor is formed by combining molecular fingerprints representing drug information. Finally, the FwRF is applied to implement on *Enzyme*, *Ion Channel*, *GPCR*, and *Nuclear Receptor* data sets. The results of the five-fold cross-validation experiment show that the prediction accuracy of this approach reaches 91.68%, 88.11%, 84.72% and 78.33% on four benchmark data sets, respectively. To further validate the performance of the RFDTI, we compare it with other excellent methods and Support Vector Machine (SVM) model. In addition, 7 of the 10 highest predictive scores in predicting novel DTIs were validated by relevant databases. The experimental results of cross-validation indicated that RFDTI is feasible in predicting the relationship among drugs and target, and can provide help for the discovery of new candidate drugs.

## Introduction

Identifying the interaction between drugs and targets is a crux area in genomic drug discovery, which not only helps to understand various biological processes, but also contributes to the development of new drugs [1,2]. The emergence of molecular medicine and the completion of the Human Genome Project provide better conditions for the identification of new drug target proteins. Although the researchers have made a lot of efforts, only a small number of candidate drugs can be approved by the Food and Drug Administration (FDA) to enter the market so far [3–5]. An important reason for this situation is due to the inherent defects of the experimental methods. As is known to all, biological laboratory methods to identify DTIs are usually expensive, time-consuming, and are limited to small-scale studies. *In silico* methods can narrow the scope of candidate targets and provide supporting evidence for the drug target experiments, thus speeding up drug discovery. Therefore, *in silico*-based methods are urgently required to improve efficiency and reduce time in identifying potential DTIs across the genome. [6–8].

In recent years, researchers have developed a variety of *in silico*-based methods to analyze and predict DTIs [9–11]. For example, Wu *et al.* [12] proposed the SDTBNI model in 2016, which searches for unknown DTIs through new chemical entity-substructure linkages, drug-substructure linkages and known DTIs networks. Zhang *et al.* [13] proposed a novel DTIs prediction model based on LPLNI. The model uses data points reconstructed from neighborhood to calculate the linear neighborhood similarity of drug-drug. Based on biomedical related data and Linked Tripartite Network (LTN), Zong *et al.* [14] used the target-target and drug-drug similarities calculated by DeepWalk to predict DTIs. In addition, Peng *et al.* [15] combines the biological information of targets and drugs with PCA-based convex optimization algorithms to predict new DTIs using semi-supervised inference method. Ezzat *et al.* [16] used ensemble learning algorithm to predict DTIs by decrease features with subinterval features through three dimensionality reduction models. Generally speaking, drugs with chemical similarity also have similar biochemical activity, that is, they can bind to similar target proteins. Based on the above assumptions, the use of medicinal chemical molecular structure information and protein sequence information to predict the DTIs model has achieved good results. For example, Wen *et al.* [17] extracted drug and target features from their chemical substructure and sequence information, and used deep belief network (DBN) to predict potential DTIs.

In this article, according to the assumption that the interaction between drugs and target proteins largely depend on the information of target protein sequences and drug molecular sub-structure fingerprints, a novel *in silico*-based model is proposed to infer potential DTIs. Our feature combines the fingerprint of the drug molecule structure and the protein sequence encoded by a feature extraction method called Pseudo Position-Specific Score Matrix (PsePSSM). In the experiment, we adopt the FwRF classifier to predict the results on the four DTIs benchmark data sets, including *Enzyme, Ion Channel, GPCR* and *Nuclear Receptor*. In order to verify the performance of the proposed model, we compared with SVM classifier model, different feature extraction models and existing excellent methods. Furthermore, in the case study, 7 of the top 10 DTIs with the highest prediction score of RFDTI model were confirmed. The promoting experimental results show that RFDTI has excellent performance and can effectively predict potential DTIs.

## Results and Discussion

### Evaluation Criteria

In this paper, accuracy (Accu.), sensitivity (Sen.), precision (Prec.), and Matthews correlation coefficient (MCC) are used to estimate the performance of RFDTI. Their formulas are as follows:

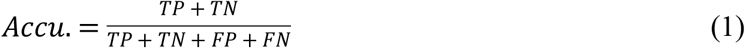

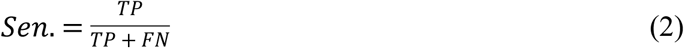

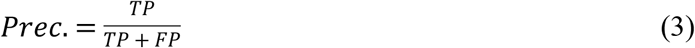

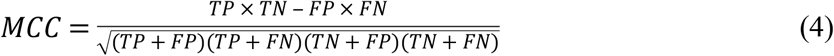

where TP is the number of drug-target pairs that are related to each other to be correctly identified; FP is the number of drug-target pairs that are related to each other to be incorrectly identified; TN is the number of drug-target pairs that are not related to each other to be correctly identified; FN is the number of drug-target pairs that are not related to each other to be incorrectly identified. Moreover, the receiver operating characteristic (ROC) curve [18–20] and area under the ROC curve (AUC) are used to visually display the performance of the classifier.

### Model Construction

To optimize the performance of the RFDTI, the grid search method is applied to explore the parameters of PsePSSM and FwRF. In the experiment we explored the effects of different PsePSSM parameters on the performance of classifiers. After optimization, we set the parameter λ of PsePSSM to 34, and the parameters the feature selection ratio *r*, the feature subset *K* and the decision tree number *L* of FwRF classifier to 0.8, 16 and 21, respectively. Figure 1 display the prediction results of different FwRF parameters, where an optimal choice of K=16 and L=21 are finally selected.

**Fig 1.**
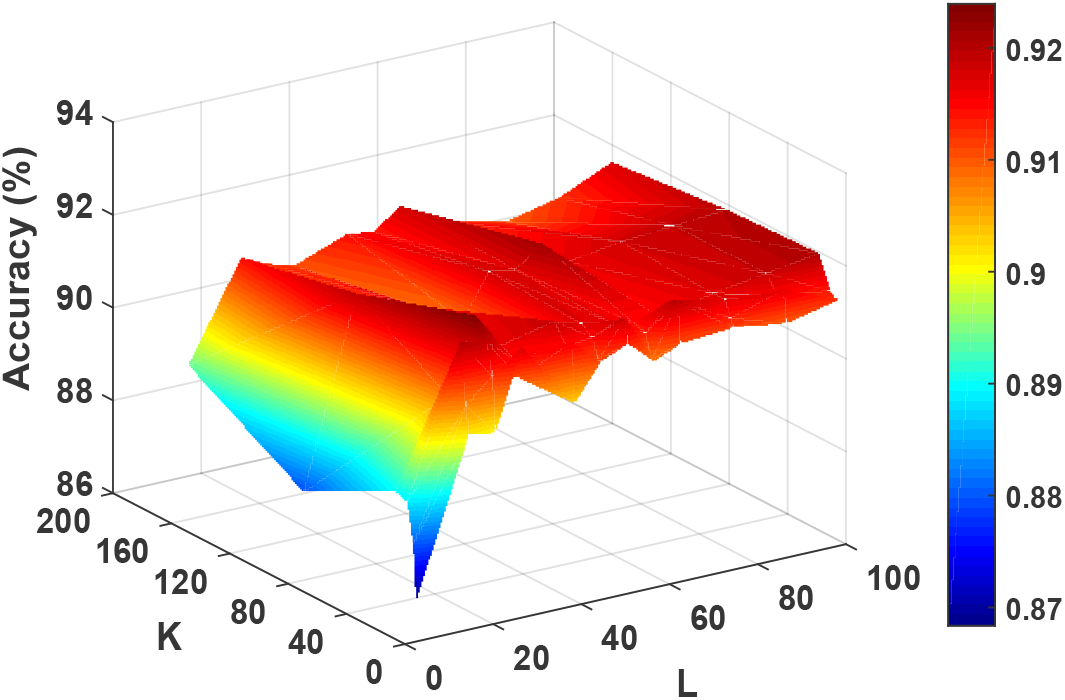
The accuracy generated by FwRF under different parameters K and L

### Evaluation of Model Prediction Ability

After finding the optimal parameters of the RFDTI, we put them in benchmark data sets, including *Enzyme*, *Ion Channel*, *GPCR* and *Nuclear Receptor.* In order to avoid over-fitting of the model, we use five-fold cross-validation method to evaluate the performance of the model. More specifically, we split the data set into five subsets, one of which is taken as the test set, and the remaining four are used as the training set. Then, the cross-validation process will be repeated five rounds. The results from the 5 times are then averaged to produce the final result.

Tables 1–4 list the predicted results by RFDTI on four benchmark data sets. In *Enzyme* data set, we gained the average of accuracy, sensitivity, precision, MCC, and AUC were 91.68%, 90.84%, 92.39%, 83.39%, and 91.72%. Their standard deviations were 0.84%, 1.68%, 1.37%, 1.68%, and 1.06%. In *Ion Channel* data set, we achieved these evaluation criteria were 88.11%, 90.30%, 86.57%, 79.02%, and 88.27%. Their standard deviations were 1.01%, 1.61%, 2.29%, 1.55%, and 1.36%. In *GPCR* data set, we yielded the average of these evaluation criteria were 84.72%, 84.73%, 84.73%, 74.06%, and 85.57%. Their standard deviations were 1.94%, 3.45%, 4.21%, 2.68%, and 2.28%. In *Nuclear Receptor* data set, we gained the average of these evaluation criteria were 78.33%, 81.97%, 78.08%, 65.56%, and 75.31%. Their standard deviations were 5.34%, 7.85%, 12.56%, 6.05%, and 5.87%. Figures 2–5 draw the ROC curve generated from RFDTI on the four benchmark data sets.

**Table 1.**
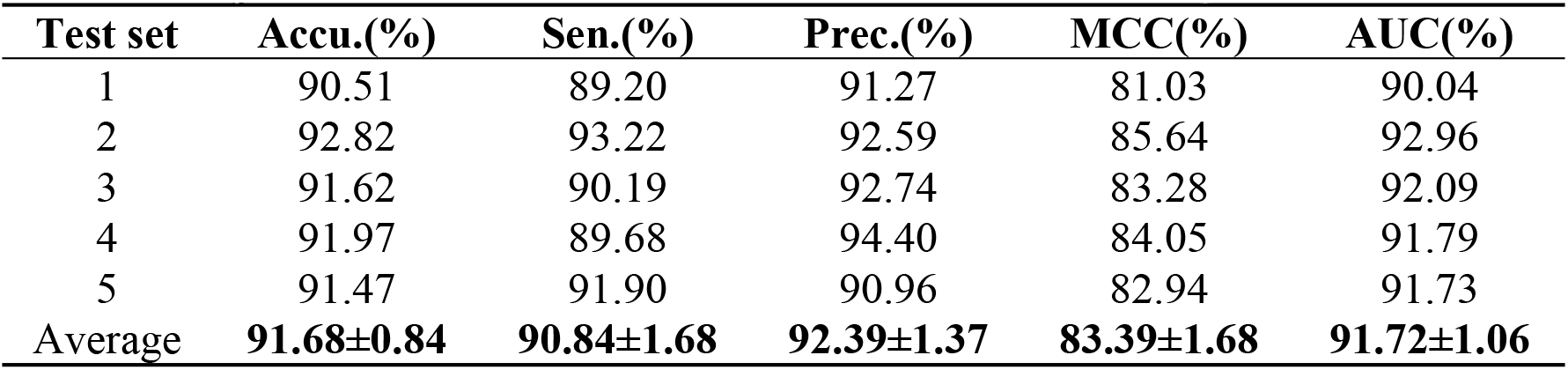
Experimental results of cross-validation of RFDTI on *Enzyme* data set.

| Test set | Accu.(%) | Sen.(%) | Prec.(%) | MCC(%) | AUC(%) |
| --- | --- | --- | --- | --- | --- |
| 1 | 90.51 | 89.20 | 91.27 | 81.03 | 90.04 |
| 2 | 92.82 | 93.22 | 92.59 | 85.64 | 92.96 |
| 3 | 91.62 | 90.19 | 92.74 | 83.28 | 92.09 |
| 4 | 91.97 | 89.68 | 94.40 | 84.05 | 91.79 |
| 5 | 91.47 | 91.90 | 90.96 | 82.94 | 91.73 |
| Average | **91.68±0.84** | **90.84±1.68** | **92.39±1.37** | **83.39±1.68** | **91.72±1.06** |

**Table 2.** Experimental results of cross-validation of RFDTI on *Ion Channel* data set.

| Test set | Accu.(%) | Sen.(%) | Prec.(%) | MCC(%) | AUC(%) |
| --- | --- | --- | --- | --- | --- |
| 1 | 86.61 | 90.38 | 83.76 | 76.76 | 86.16 |
| 2 | 87.63 | 91.92 | 84.78 | 78.22 | 87.83 |
| 3 | 88.98 | 91.61 | 87.22 | 80.36 | 89.68 |
| 4 | 88.31 | 89.67 | 87.62 | 79.33 | 89.07 |
| 5 | 89.02 | 87.93 | 89.47 | 80.43 | 88.59 |
| Average | **88.11±1.01** | **90.30±1.61** | **86.57±2.29** | **79.02±1.55** | **88.27±1.36** |

**Table 3.** Experimental results of cross-validation of RFDTI on *GPCR* data set.

| Test set | Accu.(%) | Sen.(%) | Prec.(%) | MCC(%) | AUC(%) |
| --- | --- | --- | --- | --- | --- |
| 1 | 82.28 | 86.21 | 77.52 | 70.73 | 82.86 |
| 2 | 87.01 | 88.62 | 85.16 | 77.38 | 88.82 |
| 3 | 86.22 | 86.52 | 88.41 | 76.00 | 86.72 |
| 4 | 84.63 | 80.33 | 86.73 | 73.83 | 84.37 |
| 5 | 83.46 | 81.95 | 85.83 | 72.37 | 85.11 |
| Average | **84.72±1.94** | **84.73±3.45** | **84.73±4.21** | **74.06±2.68** | **85.57±2.28** |

**Table 4.** Experimental results of cross-validation of RFDTI on *Nuclear Receptor* data set.

| Test set | Accu.(%) | Sen.(%) | Prec.(%) | MCC(%) | AUC(%) |
| --- | --- | --- | --- | --- | --- |
| 1 | 69.44 | 83.33 | 65.22 | 55.90 | 72.22 |
| 2 | 77.78 | 85.00 | 77.27 | 64.34 | 69.69 |
| 3 | 80.56 | 92.31 | 66.67 | 67.47 | 74.25 |
| 4 | 83.33 | 77.78 | 87.50 | 72.05 | 75.31 |
| 5 | 80.56 | 71.43 | 93.75 | 68.03 | 85.08 |
| Average | **78.33±5.34** | **81.97±7.85** | **78.08±12.56** | **65.56±6.05** | **75.31±5.87** |

**Fig 2.**
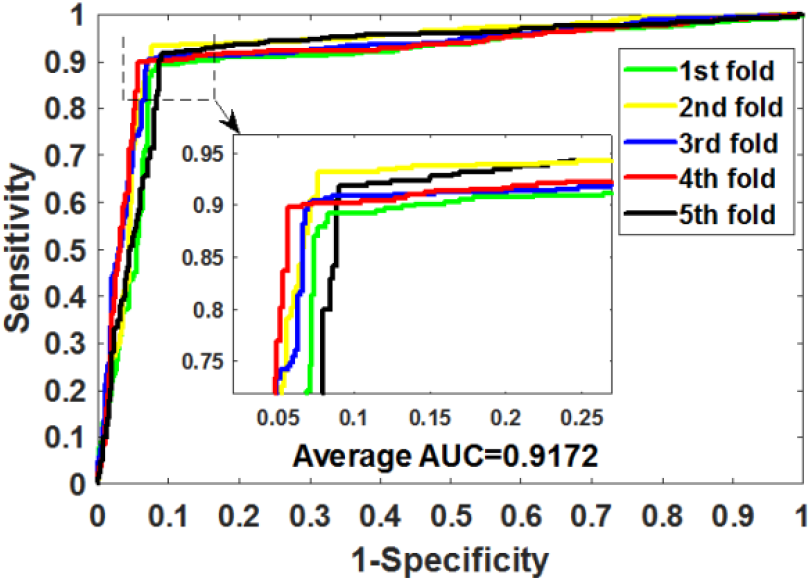
ROC curves obtained by RFDTI on *Enzyme* data set.

**Fig 3.**
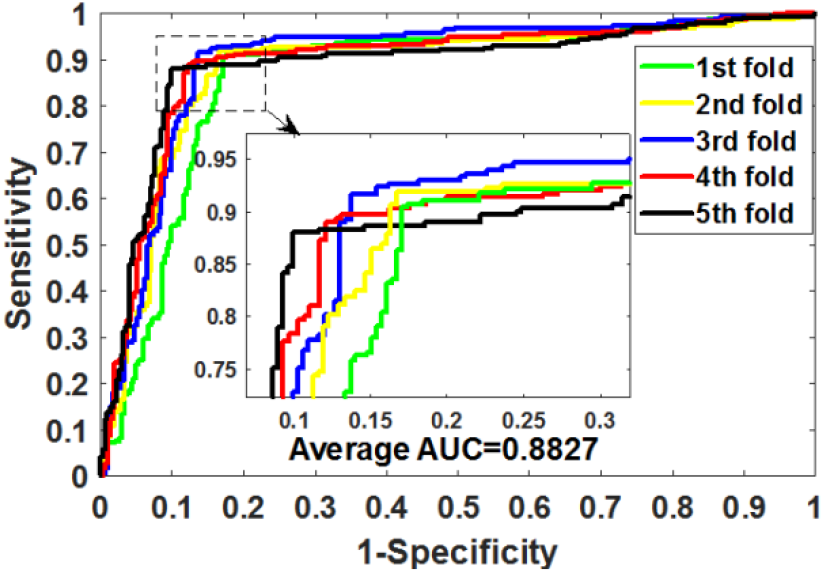
ROC curves obtained by RFDTI on *Ion Channel* data set.

**Fig 4.**
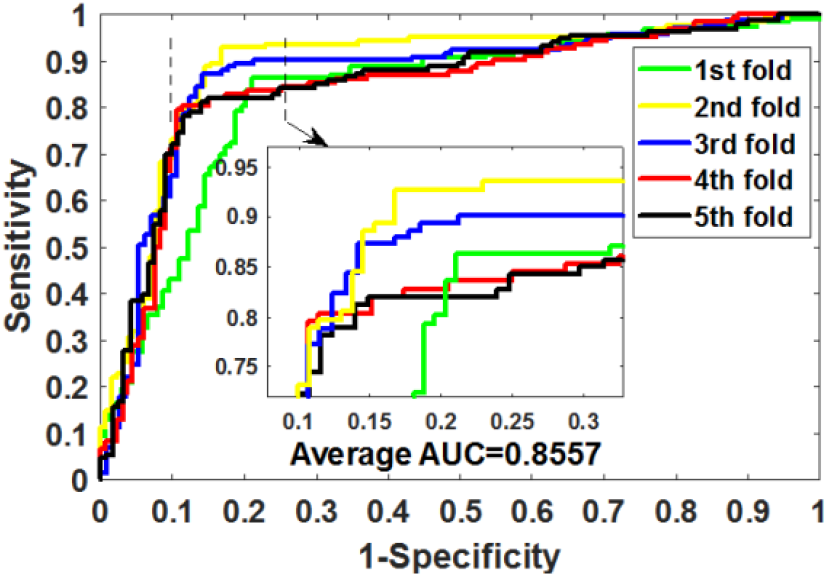
ROC curves obtained by RFDTI on *GPCR* data set.

**Fig 5.**
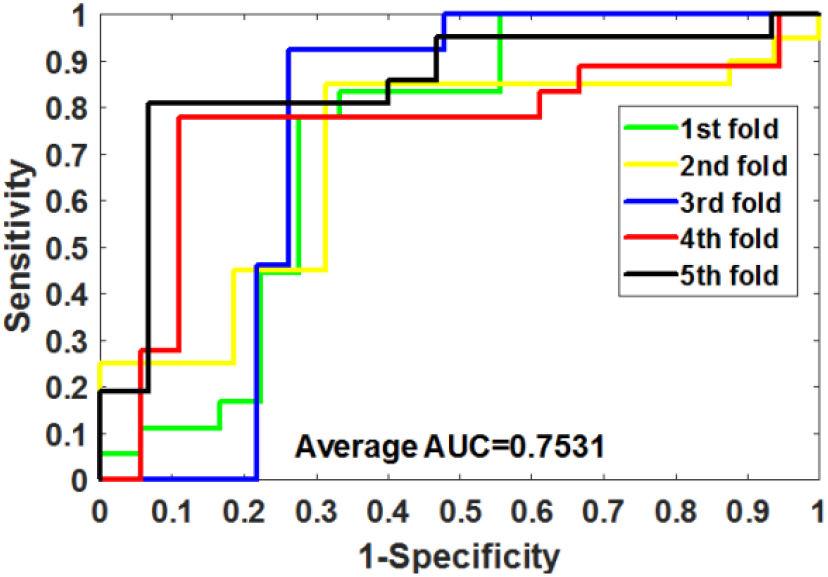
ROC curves obtained by RFDTI on *Nuclear Receptor* data set.

**Fig 6.**
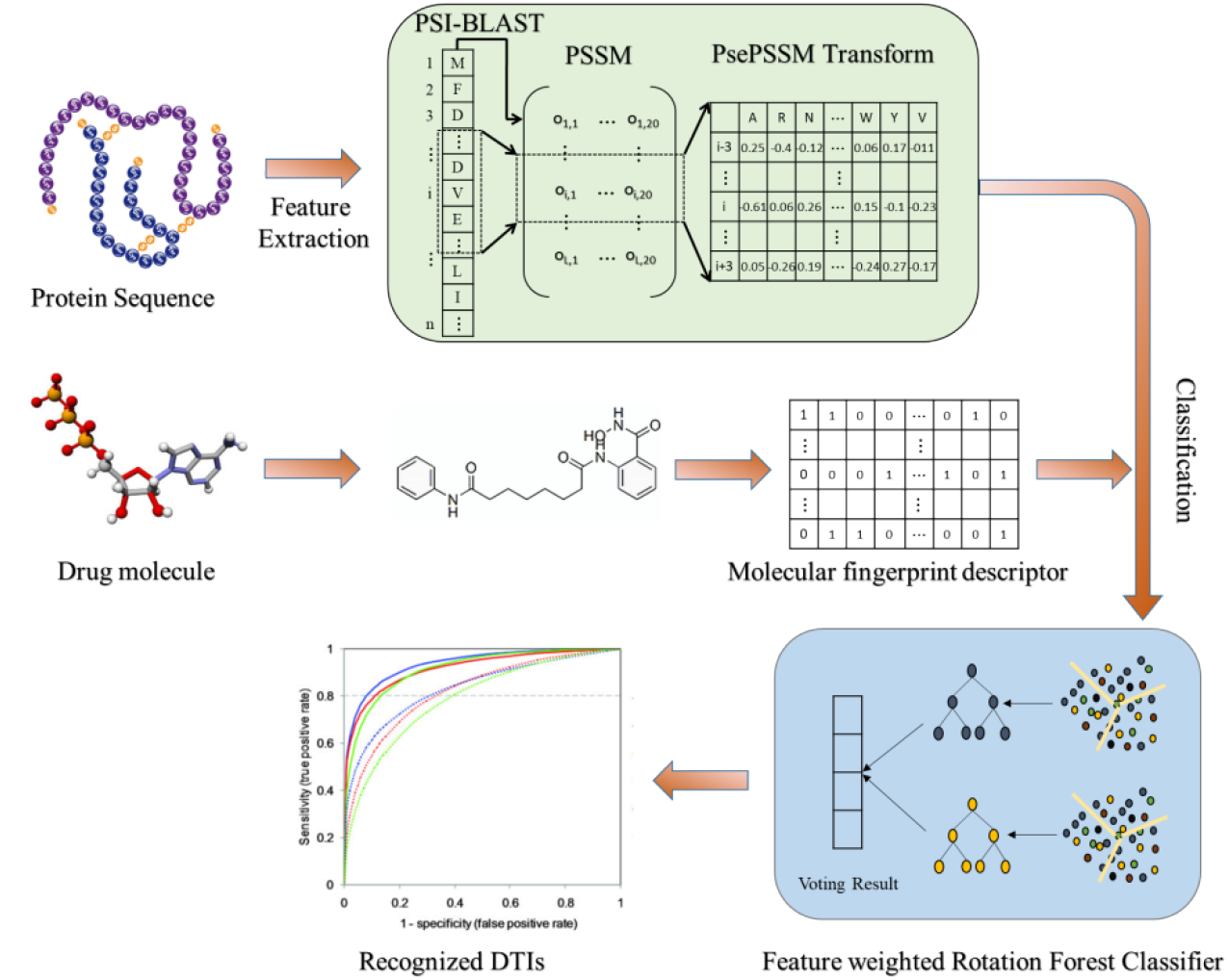
The flow chart of the proposed model

### Comparison between RFDTI and LPQ descriptor model

To evaluate the impact of PsePSSM algorithm on the proposed model, we compare it with Local Phase Quantization (LPQ) on four benchmark data sets in this section. The LPQ feature extraction algorithm is based on the blur invariance property of the Fourier phase spectrum and originally described in the article for texture description by Ojansivu and Heikkila [21]. Table 5 summarizes the cross-validation results generated by LPQ algorithm combined with FwRF classifier on four benchmark data sets. From the table we can see that RFDTI has achieved the best results in all the evaluation indicators including accuracy, sensitivity, precision, MCC and AUC. Detailed five-fold cross-validation results on four benchmark data sets are presented in Supplementary Materials Tables S1-S4. In the comparison experiment, we set the same parameters for the FwRF classifier. We can see from the comparison results that PsePSSM algorithm combined with FwRF classifier does helps to improve the performance of the model.

**Table 5.** Results of comparison experiments between RFDTI and LPQ descriptor model on four benchmark data sets

| **Data set** | **Method** | **Accu.(%)** | **Sen.(%)** | **Prec.(%)** | **MCC(%)** | **AUC(%)** |
| --- | --- | --- | --- | --- | --- | --- |
| *Enzyme* | FwRF+LPQ | 89.63±0.39 | 89.69±1.82 | 89.64±2.16 | 79.32±0.79 | 89.40±0.98 |
|  | RFDTI | **91.68±0.84** | **90.84±1.68** | **92.39±1.37** | **83.39±1.68** | **91.72±1.06** |
| *Ion Channel* | FwRF+LPQ | 83.97±2.32 | 86.93±3.03 | 81.89±3.66 | 68.13±4.54 | 84.66±2.01 |
|  | RFDTI | **88.11±1.01** | **90.30±1.61** | **86.57±2.29** | **79.02±1.55** | **88.27±1.36** |
| *GPCR* | FwRF+LPQ | 82.52±2.17 | 83.87±3.58 | 81.79±3.78 | 65.19±4.15 | 83.19±1.79 |
|  | RFDTI | **84.72±1.94** | **84.73±3.45** | **84.73±4.21** | **74.06±2.68** | **85.57±2.28** |
| *Nuclear Receptor* | FwRF+LPQ | 66.67±7.35 | 67.64±16.23 | 67.97±9.98 | 35.46±10.89 | 69.56±6.85 |
|  | RFDTI | **78.33±5.34** | **81.97±7.85** | **78.08±12.56** | **65.56±6.05** | **75.31±5.87** |

### Comparison between RFDTI and SVM classifier model

As the most versatile Support Vector Machine (SVM) classifier has been widely used by various problems. In order to estimate RFDTI clearly, we compare the results of RFDTI and SVM classifier model on the same data set. The SVM parameters are determined by grid search, and finally set the value of *c* to 0.5 and the value of *g* to 0.6. The results of the SVM classifier optimization can be viewed in the Supplementary Materials Table S5. From the table 6 we can see that RFDTI has achieved excellent results on the four benchmark data sets. Among the evaluation parameters accuracy, sensitivity, MCC and AUC, RFDTI have achieved the highest results, and RFDTI on precision is only slightly lower than that of SVM classifier model in *Enzyme* and *Ion Channel* data sets. Detailed five-fold cross-validation results on four benchmark data sets are presented in Supplementary Materials Tables S6-S9. This result indicates that the FwRF classifier is suitable for the proposed model and can effectively improve the performance of the model.

**Table 6.** Results of comparison experiments between RFDTI and SVM classifier model on four benchmark data sets

| Data set | Method | Accu.(%) | Sen.(%) | Prec.(%) | MCC(%) | AUC(%) |
| --- | --- | --- | --- | --- | --- | --- |
| *Enzyme* | PsePSSM+SVM | 84.20±0.60 | 69.90±1.70 | **98.00±0.50** | 71.50±1.00 | 84.30±1.20 |
|  | RFDTI | **91.68±0.84** | **90.84±1.68** | 92.39±1.37 | **83.39±1.68** | **91.72±1.06** |
| *Ion Channel* | PsePSSM+SVM | 81.90±1.20 | 69.70±3.70 | **92.40±2.20** | 66.00±1.90 | 81.70±1.20 |
|  | RFDTI | **88.11±1.01** | **90.30±1.61** | 86.57±2.29 | **79.02±1.55** | **88.27±1.36** |
| *GPCR* | PsePSSM+SVM | 70.00±2.10 | 50.40±7.80 | 82.30±3.30 | 42.80±4.90 | 70.10±2.70 |
|  | RFDTI | **84.72±1.94** | **84.73±3.45** | **84.73±4.21** | **74.06±2.68** | **85.57±2.28** |
| *Nuclear Receptor* | PsePSSM+SVM | 63.30±3.60 | 57.60±7.90 | 67.50±14.60 | 29.60±7.40 | 61.80±5.80 |
|  | RFDTI | **78.33±5.34** | **81.97±7.85** | **78.08±12.56** | **65.56±6.05** | **75.31±5.87** |

### Comparison with existing methods

The prediction of the relationship between drugs and targets has drawn increasing interest of researchers. So far, a lot of excellent computational approaches have been designed. To better verify the proposed approach, we compare it with other existing methods using five-fold cross-validation on the same benchmark data sets. Table 7 lists the details of other excellent methods and RFDTI on four benchmark data sets in terms of the AUC. It can be seen that the results obtained by RFDTI on *Enzyme* and *Ion Channel* data sets are significantly higher than those of other existing methods, and the results achieved on *GPCR* data sets by RFDTI only lower than the highest result 1.13%. The performance of RFDTI on *Nuclear Receptor* data set is not very good, it may be because the sample number of the *Nuclear Receptor* data set is too small, and the training of the classifier is not sufficient

**Table 7.** Performances of other excellent methods and RFDTI on four benchmark data sets in terms of the AUC.

| Data set | RFDTI | MLCLE [10] | NetCBP [2] | SIMCOMP[22] | AM-PSSM[4] |
| --- | --- | --- | --- | --- | --- |
| *Enzyme* | 0.9172 | 0.842 | 0.8251 | 0.863 | 0.843 |
| *Ion Channel* | 0.8827 | 0.795 | 0.8034 | 0.776 | 0.722 |
| *GPCR* | 0.8557 | 0.850 | 0.8235 | 0.867 | 0.839 |
| *Nuclear Receptor* | 0.7531 | 0.790 | 0.8394 | 0.856 | 0.767 |

### Case study

To further validate RFDTI’s ability to predict potential DTIs, we use all known interactions to train the model and then predict unknown interactions. We selected 10 drug-target pairs with the highest predictive score to validate in SuperTarget [23]. SuperTarget is a database that collects drug-target relations and currently stores 332,828 DTIs. As shown in Table 8, 7 of the top 10 predicted highest scores were confirmed by the proposed model. This result indicates that RFDTI can effectively predict the potential DTIs. It is worth noting that although we have not found evidence of the interaction of the remaining 3 drug-target pairs, we cannot completely deny the possibility of their interactions.

**Table 8.**
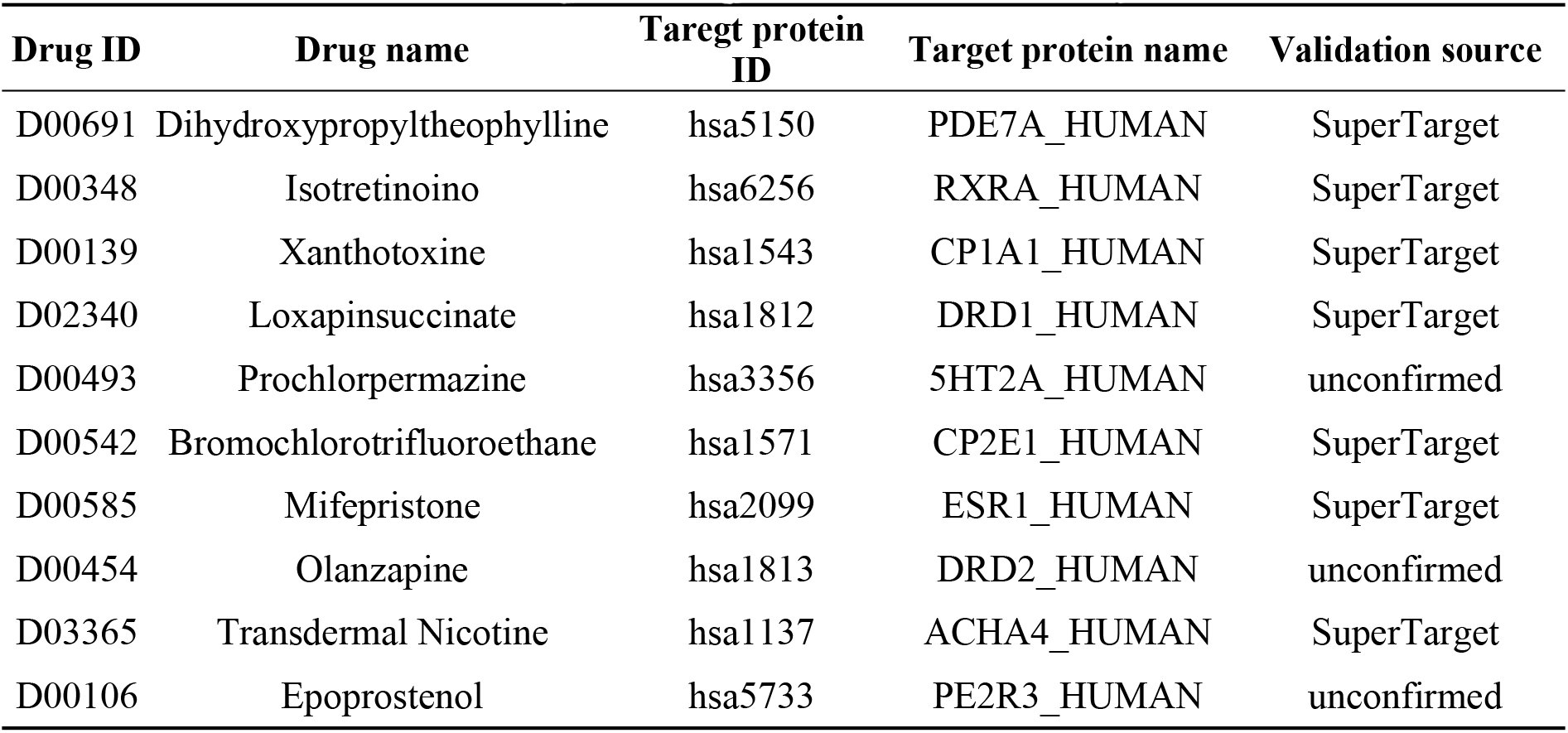
The top 10 new predicted interactions by RFDTI

| Drug ID | Drug name | Taregt protein ID | Target protein name | Validation source |
| --- | --- | --- | --- | --- |
| D00691 | Dihydroxypropyltheophylline | hsa5150 | PDE7A_HUMAN | SuperTarget |
| D00348 | Isotretinoino | hsa6256 | RXRA_HUMAN | SuperTarget |
| D00139 | Xanthotoxine | hsa1543 | CP1A1_HUMAN | SuperTarget |
| D02340 | Loxapinsuccinate | hsa1812 | DRD1_HUMAN | SuperTarget |
| D00493 | Prochlorperazine | hsa3356 | 5HT2A_HUMAN | unconfirmed |
| D00542 | Bromochlorotrifluoroethane | hsa1571 | CP2E1_HUMAN | SuperTarget |
| D00585 | Mifepristone | hsa2099 | ESR1_HUMAN | SuperTarget |
| D00454 | Olanzapine | hsa1813 | DRD2_HUMAN | unconfirmed |
| D03365 | Transdermal Nicotine | hsa1137 | ACHA4_HUMAN | SuperTarget |
| D00106 | Epoprostenol | hsa5733 | PE2R3_HUMAN | unconfirmed |

## Materials and Methodology

### Benchmark data sets

In this article, we applied four protein targeting data sets, including *Enzyme, Ion Channel, GPCR* and *Nuclear Receptor*. These data sets are applied as the benchmark data sets by Yamanishi *et al.* [24] and collected from the BRENDA, DrugBank, SuperTarget & Matador and KEGG BRITE. They can be downloaded at http://web.kuicr.kyoto-u.ac.jp/supp/yoshi/drugtarget/. The number of drugs was 445, 210, 233 and 54, and the number of target proteins was 664, 204, 95 and 26 in these benchmark data sets, respectively. Among these data, 5127 pairs of drug-target were confirmed to interact with each other, corresponding to 2926, 1476, 635 and 90 pairs in four data sets, respectively [25].

The DTIs network can be expressed by a bipartite graph in which nodes represent drugs or targets, and edges represent their interactions. If there is a relationship between the nodes, connect them with edges, otherwise they do not connect. The edges in the initial bipartite graph represent the real DTIs have been detected by biological experiments. The number of initial edges is relatively small compared to a completely connected bipartite graph [26–29]. For example, there are totally 54×26=1404 connections in the bipartite graph of the *Nuclear Receptor* data set. However, the experiment detected representatives known drug-target interactions only 90. Therefore, the number of positive drug-target pairs (e.g., 90) accounted for only 6.41% of the total number of drug-target pairs (e.g., 1404), much less than the number of negative drug-target pairs (e.g., 1404-90=1314). The same problem also appears in the other three data sets. In order to solve the problem of data imbalance, we randomly select negative drug-target pairs with the same number of positive drug-target pairs. In fact, such negative samples may contain drug-target pairs with interactions. To reduce this possibility, we randomly selected ten times negative sample sets in the experiment. From a statistical point of view, the number of real interactions on a large bipartite graph is chosen as the negative sample set is very small. Eventually, the negative sample numbers of *Enzyme, Ion Channel, GPCR* and *Nuclear Receptor* data sets are 2926, 1476, 635 and 90, respectively.

### Molecules description

In recent years, different types of descriptors have been proposed to represent drug compounds, such as quantum chemical properties, topological, constitutional and geometrical. Since the molecular substructure fingerprint does not require the three-dimensional structural information of the molecule and has the advantage of directly reflecting the relationship between molecular properties and structure, more and more researchers use it as a descriptor to predict the relationship between the drug and the target protein. Specifically, we first store all the molecular substructures in the form of a dictionary, and then split a given drug molecule. When it contains a certain substructure, the corresponding bit of the descriptor is assigned to 1; otherwise it is assigned to 0. Finally, we get the drug molecule in the form of Boolean vectors. In the experiment, we use the chemical structure fingerprint set from PubChem System, and the fingerprints property is “PUBCHEM_CACTVS_SUBGRAPHKEYS” in PubChem. A drug fingerprint is recorded as 881 substructures, so the drug molecule feature is the 881-dimensional. Since the drug fingerprint is divided into 881 substructures, the dimension of the drug molecular fingerprint descriptor is 881 dimensions.

### Numerical characterization of protein sequences

In the experiment, we used Position-Specific Scoring Matrix (PSSM) to convert protein sequence numerically [30]. PSSM is widely used in protein binding site prediction, protein secondary structure prediction, and prediction of disordered regions. PSSM is an *L* × *20* matrix that can be expressed as *S* = {∂_*i*,*j*_:*i* = 1⋯*L* and *j* = 1⋯20}, where 20 represents the number of the amino acids and *L* denotes the length of the protein sequence. PSSM matrix *S*(*i*,*j*) can be expressed as follows:

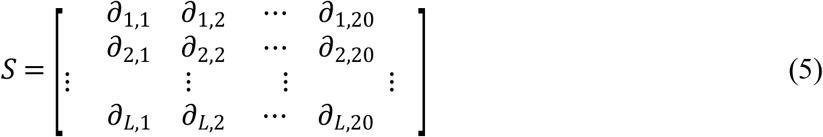

Where ∂_*i*,*j*_ denotes the probability that the ith residue being mutated into the jth amino acid during the evolutionary process of protein multiple sequence alignment.

In the experiment, we use Position-Specific Iterated BLAST (PSI-BLAST) tool [31] to generate PSSM based on *SwansProt* data set. PSI-BLAST can generates the 20-dimensional vector indicating the mutation conservation probabilities of 20 different amino acids. To obtain high homologous and broad homologous sequences, we set the parameter iterations to 3 and the parameter e-value to 0.001, and keep other parameters as default values. The *SwissProt* database and PSI-BLAST toolkit can be downloaded at http://blast.ncbi.nlm.nih.gov/Blast.cgi.

### Feature extraction algorithm

Effective protein feature descriptors can not only mine useful information, but also improve the performance of the approach. In this study, we introduce the feature extraction algorithm Pseudo Position-Specific Score Matrix (PsePSSM), which concept from Chou *et al.* [32]. The PsePSSM is expressed by formula as follows:

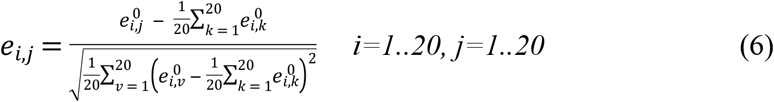

where 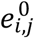 denotes the raw score generated by PSI-BLAST, which is typically a positive or negative integer. This is not the final score, because if it exceeds 20 amino acids, the score may contain 0; if the same conversion procedure continues, the score may remain unchanged. The positive number signifies that the frequency of corresponding mutations in the alignment is higher than that of accidental expectations. Conversely, the negative number signifies that the frequency of corresponding mutations in the alignment is lower than that of accidental expectations. However, based on the PSSM formula, proteins of different lengths will produce a matrix of different numbers of rows. Therefore, equation 7 is used to convert the PSSM matrix into a uniform pattern.

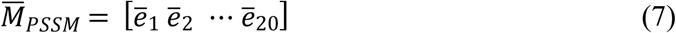

and

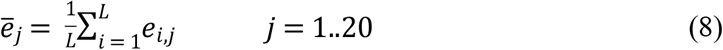

where *e̅*_*j*_ indicates the average score of *P* protein when amino acid residues evolve into j-type amino acids. However, if only *M̅*_*PSSM*_ is used to indicate protein *P*, all information about sequence order will be lost during evolution. In order to prevent this from happening, we introduce the idea of pseudo amino acid to improve equation 7 Therefore, according to the formula 9, we can get the features of segmented PsePSSM:

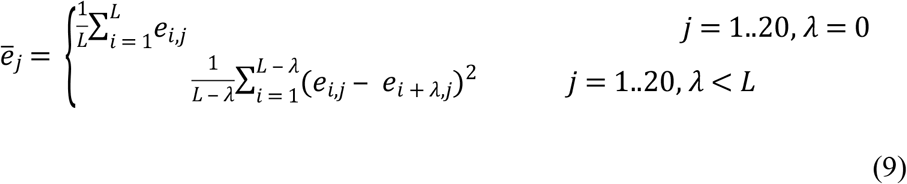

where *e*_*j*_ is a related factor for j-type amino acid, whose contiguous distance is along each segmented protein sequence. The flow chart of the proposed model is shown below.

### Feature Weighted Rotation Forest Classifier

In this paper, we use the feature weighted rotation forest (FwRF) [33–35] to accurately predict DTIs. Compared with the original rotation forest, FwRF adds the function of weight selection. Through FwRF we can remove the noise features with small weights, thus increasing the content of useful information and improving the accuracy of prediction. The weights of the features are calculated by *χ*^2^ statistical method. The feature *F* for the class can be obtained by the following formula.

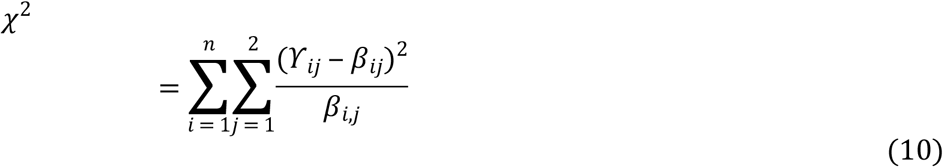

where *n* is the number of values in feature *F*, *ϒ*_*ij*_ is the number of features in the *f*_*j*_ class with a value of *v*_*i*_, which can be expressed as a formula:

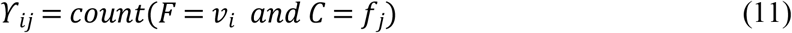

*β*_*i*,*j*_ is the expected value of *v*_*i*_ and *f*_*j*_ which is defined as follows:

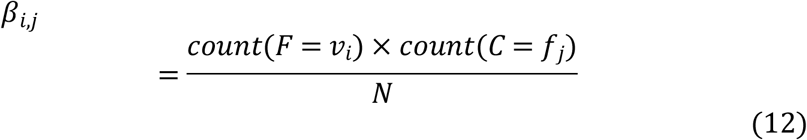

where *count*(*C* = *f*_*j*_) is the number of samples with the value of *f*_*j*_ in class *C*, *count*(*F* = *v*_*i*_) is the number of samples whose value of feature *F* is *v*_*i*_, and *N* is the number of samples.

The implementation steps of feature weighted rotation forest are as follows: Firstly, the weights of all features are calculated by equation 10; secondly, the features are sorted according to the weights; finally, the desired features are selected according to the given feature selection rate *r*. After performing these steps, we get a new data set and send it to rotation forest.

Assuming {*x*_*i*_,*y*_*i*_} contains *S* training samples, where in *x*_*i*_ = (*x*_*i*1_,*x*_*i*2_,…,*x*_*in*_) be an n-dimensional feature vector. Let *X* is the training sample set, *Y* is the corresponding labels and *F* is the feature set. Then *X* is *S*×*n* matrix, which is composed of n observation feature vector composition. Assuming that the number of decision trees is *N*, then the decision trees can be expressed as *D*_1_,*D*_2_,…,*D*_*N*_. The algorithm is executed in the following steps.

1. Using the appropriate parameter K to randomly divide *F* into *K* independent and uncrossed subsets, the number of each subset feature is 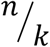.
2. A corresponding column of features in the subset *D*_*i*,*j*_ is selected from the training set *X* to form a new matrix *X*_*i*,*j*_. Then, 75% of the data is extracted from *X* in the form of bootstrap to form a new set *X′*_*i*,*j*_.
3. Use matrix *X′*_*i*,*j*_ as the feature transform to generate coefficients in matrix *M*_*i*,*j*_.
4. Using the coefficients obtained from the matrix *M*_*i*,*j*_ to form a sparse rotation matrix *R*_*i*_, the expression of which is as follows:

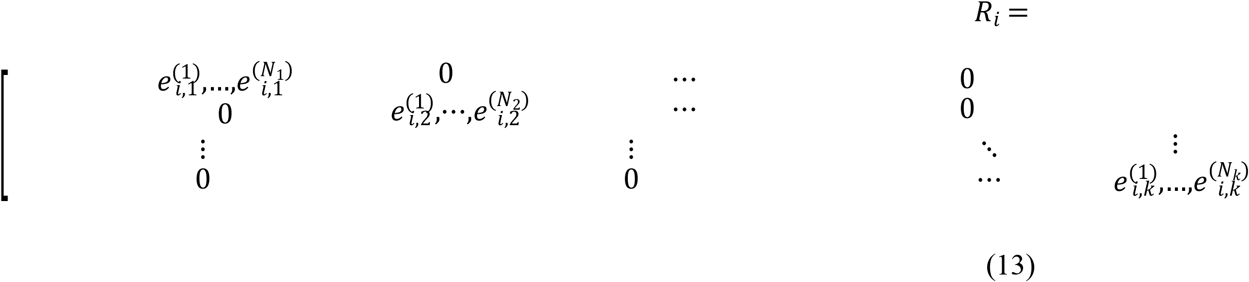

In classification, the test sample *x* is determined to belong to the *y*_*i*_ class by the *x* generated by the classifier *D*_*i*_ of 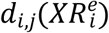. Then calculate the confidence class by the following average combination formula:

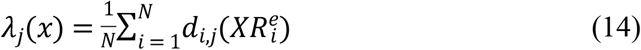

Finally, the class with the largest value *λ*_*j*_(*x*) is discriminated as *x*.

### Conclusions

Prediction of DTIs is a crucial problem for human medical improvement and genomic drug discovery. Under the hypothesis that the drug molecules structures and protein amino acids sequence have a big impact on the relationships among drugs and target proteins, the RFDTI model is proposed to infer potential drug-target relationships in this article. We implement it on *Enzyme*, *Ion Channel, GPCR* and *Nuclear Receptor* data sets, and obtained excellent results. To further evaluate the performance of the proposed approach, we compared it with PsePSSM model, the SVM classifier model and other existing methods on the same data sets. Moreover, 7 of the top 10 drug-target pairs predicted by the RFDTI model were confirmed by independent data set. Competitive cross-validation experimental results show that the performance of RFDTI has been significantly improved, which demonstrated RFDTI is stable and reliable.

## Acknowledgments

This work is supported in part by the National Natural Science Foundation of China, under Grants 61572506, 61702444, and in part by the Pioneer Hundred Talents Program of Chinese Academy of Sciences, and in part by the CCF-Tencent Open Fund, in part by the Chinese Postdoctoral Science Foundation, under Grant 2019M653804, and in part by the West Light Foundation of The Chinese Academy of Sciences, under Grant 2018-XBQNXZ-B-008.

